# The GalNAc-T Activation (GALA) Pathway: Drivers and Markers

**DOI:** 10.1101/301978

**Authors:** Joanne Chia, Felicia Tay, Frederic Bard

## Abstract

The enzymes GALNTs add GalNAc sugar to Ser and Thr residues, forming the Tn glycan. GALNTs are activated by trafficking from Golgi to ER, a process driven by the Src kinase and negatively regulated by ERK8. This GALNTs activation (aka GALA) pathway induces high Tn levels and is a key driver of liver tumor growth. Recently, Tabak and colleagues have contested our previous data that EGF stimulation can induce GALNTs relocation. Here, we show that relocation induced by EGF is actually detectable in the images acquired by Tabak et al. Furthermore, we show that expression of EGFR enhances relocation and appears required to drive relocation induced by ERK8 depletion. We also propose that quantification of O-glycosylation of the ER resident protein PDIA4 provides an alternative measure of GALA. In sum, we demonstrate that non-reproducibility was due to experimental errors, that EGFR is indeed a driver of GALA and propose additional markers to facilitate the study of this pathway.

## Introduction

Replicability is essential to the scientific progress and has been the subject of intense debate in recent years. In biomedical sciences, some authors have argued that a large fraction of scientific studies are unreproducible, calling into question the value of discoveries and initiating a fierce debate [1–3].

In a study posted on BioRxiv and later published, Tabak and colleagues questioned the replicability of findings we published in 2010 and the physiological relevance of the GALNTs Activation (GALA) pathway [4]. In the 2010 paper, we proposed that GALNTs enzymes are regulated through trafficking from the Golgi to the ER. We showed that this relocation is induced by the tyrosine kinase Src. We further proposed that stimulation of cells by growth factors such as EGF and PDGF is able to induce this relocation, consistent with one proposed mode of activation of Src. We showed evidences that the Arf1-COPI machinery responsible for Golgi to ER traffic is involved in this relocation. Furthermore, we showed evidences that GALNTs are active in the ER and that their activity is stimulated by the relocation, constituting a potent mechanism to control O-glycosylation, which we named the GALA pathway. O-GalNAc glycosylation occurs on thousands of secreted and cell surface proteins and is essential for multicellular life [5–8]. O-glycans are built by the sequential addition of simple sugars. GALNTs initiate the sequence by adding an N-Acetylgalactosamine (GalNAc) to a Ser or Thr residue. The resulting structure is called the Tn antigen and recognised by lectins such as VVL and HPL. Tn is usually a biosynthetic intermediate that is modified by the addition of other sugars such as galactose by the enzyme C1GALT [9]. Further extension can result in more complex O-glycans. Tn can also be sialylated by sialyl-transferases, producing Sialyl-Tn, which is not further modified [10].

The formation of Tn has received a lot of attention from glycobiologists because malignant tumors have long been reported to have dramatically increased levels of Tn [11,12]. The phenotype is shared by most types of solid malignant tumors and occuring at a frequency of 70-90%, begging the question of its underlying molecular mechanisms and role in tumor biology. A proposed mechanism is that cancer cells lose the activity of the Tn modifying enzymes, in particular through mutations or loss of expression of its X-linked dedicated chaperone called Cosmc [13]. The selective advantage of short O-glycans for tumor cells is not clear, albeit they could interact with endogenous lectins such as Galectins and promote signaling events [14]. Studies have documented that short O-glycans can induce oncogenic features [15]. However, mutations of Cosmc have been reported to be extremely rare [15,16] and recent studies have shown that loss of C1GALT activity slows tumor progression in a mammary tumor model [17].

In our 2010 paper, we proposed that an alternative mechanism can also drive high Tn: the relocation of GALNTs to the ER induces an accumulation of Tn in this organelle. Since the ER tends to mesh the whole cytoplasm, the result is an increase of total cellular staining detectable by lectin-based histochemistry [4]. Importantly, we have not detected other O-glycans in the ER so far, suggesting that GalNAc is not modified by C1GALT and other extension enzymes. As the ER is much larger than the Golgi with a high content of proteins, the abundance of substrate and the lack of Tn modification can yield large increases in Tn levels when enzyme relocation is marked.

In a subsequent study, using VVL staining fluorescence imaging and quantitative analysis on tissue microarrays, we evaluated Tn levels increase to be up to 10 to 15 folds in human breast cancer tissues [18]. We showed that the pattern of Tn staining in these samples is consistent with an ER localisation [18]. Furthermore, expressing an ER-targeted form of GALNT2 or T1 in cell lines such as HeLa, MDA-MB-231 and HepG2, is sufficient to induce a 6 to 10 fold increase of total HPL staining intensity [4,18,19]. By contrast over-expressing the wild-type form of the enzyme has no or very limited effects on Tn. More recently, we have shown that relocation is also occurring in a majority of liver cancers in humans and in mice [19]. Expressing an ER-targeted GALNT1 in this tumor model strongly accelerates tumor growth. Together, these data suggest that GALNTs relocation to the ER has a selective advantage for tumor cells and is the main mechanism driving Tn increase in breast and liver tumors.

How is the relocation process controlled? An RNAi screening approach revealed that GALNTs relocation is under the control of a complex genetic network. Upon depletion of a number of genes, Tn cellular staining significantly increases. 12 different genes were validated [20]. In the case of the Ser/Thr kinase ERK8, Tn levels increase ranged from 6 to 15 folds depending on the experimental context. This study suggested that there might be many different mechanisms able to induce the relocation of GALNTs, which is highly dependent on the status of the intracellular signalling network. We found various evidences that pathways downstream of stimulation by growth factors such as EGF and PDGF are able to activate GALA [4,20]. However, stimulation by EGF and PDGF itself produces a relatively moderate relocation.

In their study, Tabak and colleagues used EGF and PDGF stimulation and GALNTs staining and reported that they could not observe a decrease GALNTs colocalisation with a Golgi marker or increase of colocalisation with an ER marker.

In the intervening months, we have gone back to our initial study and data sets, repeated several experiments and analysed raw data acquired by Herbomel et al. and shared during the process of publishing their study. We could confirm that EGF and PDGF stimulation does induce changes in the pattern of Tn and GALNT staining. We observed that these experiments are sensitive to culture conditions, in particular to FBS. We also found that, albeit moderate, the relocation was present in the images acquired by Herbomel but could not be quantified by the analytical method they chose. Finally, we show that EGFR expression enhances relocation and present a new marker to facilitate the modeling and measurement of the GALA relocation process in vitro and in vivo.

## Results

### Batch differences in FBS affect growth factor response

While we were analysing the causes of discrepancy between our results and those of Tabak’s group, we considered the possible influence of cell lines and culture conditions. HeLa cell lines are known to be highly variable [21]. However we had sent our cells to the NIH and it did not seem to improve their experiments.

We grow HeLa cells with 10% Fetal Bovine Serum (FBS) which contains various growth factors and cytokines. Before growth factor stimulation, the cells were washed twice with PBS before overnight starvation in serum-free media. Over the years, we observed that different batches of FBS can have an effect on basal levels of Tn. We tested whether, despite the washing step, two different batches of FBS could affect the experiment. With one batch of FBS (FBS1), we were able to observe an increase of 1.5 to 2 fold in Tn levels after 6h of EGF stimulation and to a lower extent with PDGF treatment (Fig 1A, 1B). However, we were surprised to find that when grown with another batch (FBS2), the effect of EGF or PDGF was minimal to sometimes undetectable (Fig 1C, 1D). Thus, contrary to expectation, washing and overnight FBS starvation does not completely erase the effect of previous growth conditions and can lead to a lack of response to EGF or PDGF. This sensitivity to culture conditions might partly explain the results reported by Herbomel et al.

**Fig 1.**
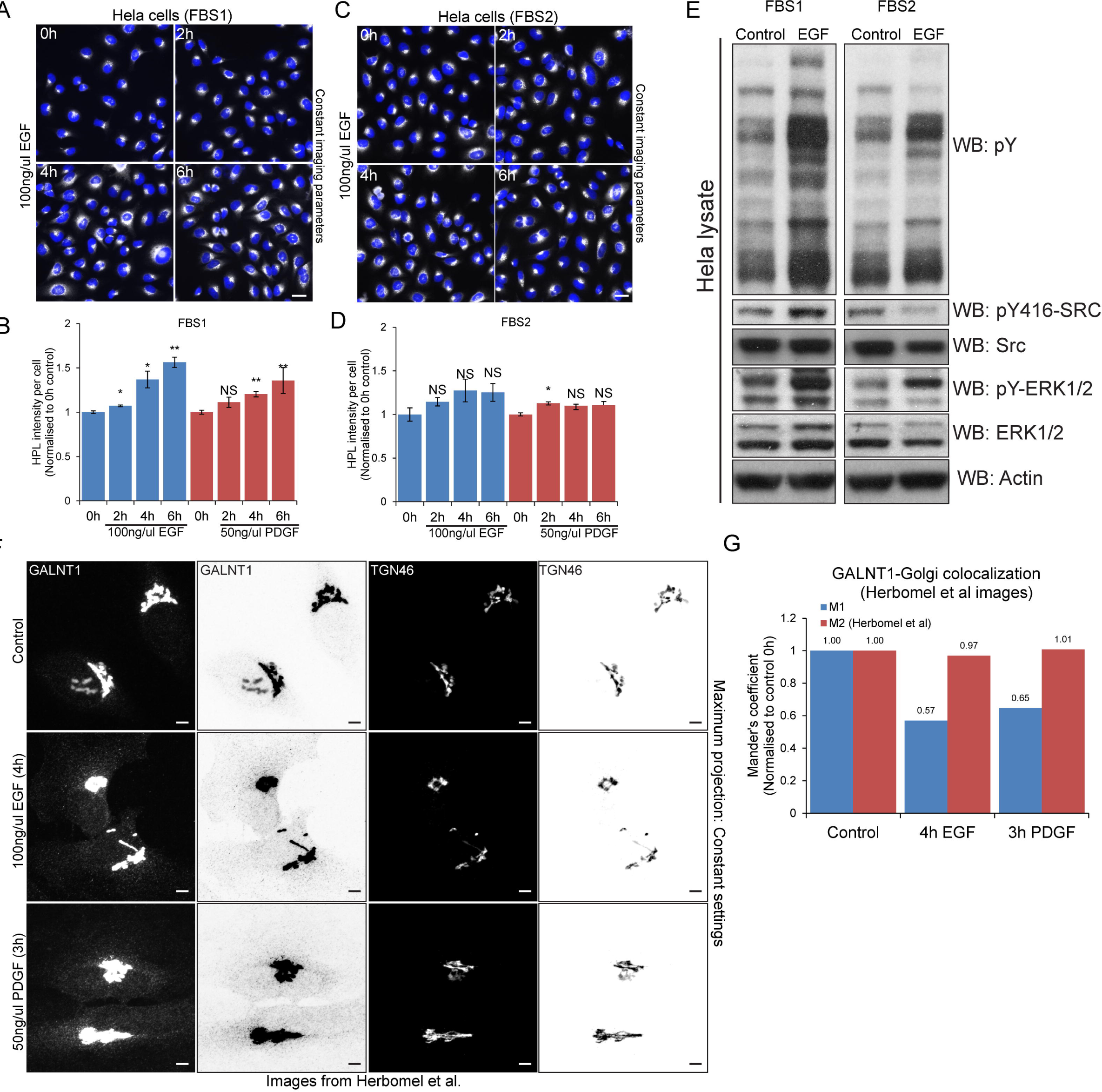
FBS type affects GALA activation while image display and analysis techniques affect interpretation of GALA activation following growth factor stimulation. *Helix pomatia* lectin (HPL) staining of Hela cells grown in (A) FBS1 and (C) FBS2 over time of EGF treatment (100 ng/ml). Images were acquired on ImageXpress Micro (IXM) at 20X magnification (air). Scale bar: 30 μm. HPL staining intensity of cells grown in (B) FBS1 and (D) FBS2 over time with 100 ng/ml EGF or 50 ng/ml PDGF stimulation. HPL intensity was analysed using the ‘Transfluor HT’ module of MetaXpress software (Molecular Devices) using the method described in [35]. Statistical significance (p) measured by two-tailed paired t test. *, p < 0.05 and **, p < 0.01 relative to mean HPL staining in unstimulated serum starved cells (0 h). NS, not significant. (E) Immunoblot analysis of Hela cells grown in FBS1 and FBS2 and were stimulated with 100 ng/ml EGF over 4 hours. Src and ERK1/2 activation and phosphotyrosine levels were analysed by western blotting (F) Maximum projection of a z-stack of 61 images from representative images provided by Herbomel et al. Each image was acquired at 60X magnification. Hela cells were stimulated with EGF and PDGF for 4 hours and 3 hours respectively. Scale bar: 5 μm. (G) Quantification of Mander’s coefficient of GALNT1 and Golgi marker TGN in (F). M1 represents the fraction of GALNT1 staining overlapping the Golgi and M2 represents the fraction of Golgi overlapping GALNT1 staining. Only M2 results were presented in Herbomel et al.

### ERK1/2 phosphorylation is not a good control for Src activation

We next repeated the same experiment to measure the levels of total phosphotyrosine and Src activation after EGF stimulation. Consistent with the results with GALA, we observed that FBS1 induced a marked activation of Src, while FBS2 had a very limited effect (Figure 1E). In their publication, Herbomel et al. used the phosphorylation of ERK1/2 as a positive control for EGF stimulation [22]. In our test, we observed that FBS2 was able to induce a clear activation of ERK1/2, even when activation of Src was undetectable (Figure 1E).

### Image analysis can significantly affect GALA quantification

Herbomel et al. used confocal imaging at high magnification with extensive sectioning (61 slices). As previously noted in our commentary of their study, this extensive sectioning can have two effects detrimental for detection of GALNTs relocation: it prevents the visualisation of weak signal and is prone to bleaching the weak signal originating from the ER [23]. Dr. Tabak had shared the images used in their publication with us. To improve the detection of weak signal, we generated a maximal intensity projection of their confocal stack then used grayscale inversion to better visualise the out-of-Golgi signal. It then became clear that in the images acquired by Herbomel, the redistribution of GALNTs is clearly visible (Figure 1E).

We next wondered why the image quantification protocol used did not detect any change. We realised that the analytical method chosen was inadequate to quantify relocation. The Manders coefficient (“M2”) used to quantify relocation measures the fraction of the Golgi marker (TGN46) colocalising with HPL or GALNT [22]. Since a significant fraction of GALNT remains at the Golgi after EGF stimulation, the fraction of TGN46 that colocalises with GALNT is unchanged. It is important to note that Manders coefficient is not sensitive to changes in staining intensities, so the M2 coefficient is not affected by a decrease in GALNT intensity. In opposition to M2, the M1 coefficient measures the fraction of GALNT that colocalises with TGN46.

We used Herbomel’s images to re-run the Manders coefficient analysis and found that indeed M2 did not register any effect; but the M1 coefficient showed a significant decrease (Fig 1G, S1C). We also applied the Manders coefficient measure on our own images and obtained similar results: no effect was detected using M2, and significant effect was detected using M1 (Figure S1A, S1B).

We could not perform the same analysis on all the images provided by Tabak’s group as some of them display inconsistent total intensity levels. There is generally lower intensities in the images of EGF stimulated cells. This is apparent in the staining of ER marker calnexin (CANX), whose levels do not change (Fig S1D, S1E). Given that Tn measures the GALNT enzyme activity levels, total Tn intensity levels are important to make comparison across treatments. We ensured that there is equal intensity levels across treatments shown in Fig 1F and found the ER marker levels to be similar in the three images (Fig S1E).

We conclude that the lack of effect reported by Herbomel et al. is due to their image analysis methods and acquisition settings.

### EGFR levels strongly affect GALA response to EGF stimulation

Our previous results collectively indicate that EGFR is a regulator of GALA. But levels of expression of this receptor are known to vary extensively; this could explain why EGF stimulation is sometimes not very effective. We thus transfected HeLa cells with EGFR, then stimulated them with EGF. This led to a marked increase in Tn levels of more than 6 fold on average (Fig 2A, 2B). The effect was relatively variable between cells, with up to 20 fold increase after 4h EGF stimulation in some cells (Fig 2B). Thus, EGF stimulation can clearly increase GALA levels and this is highly dependent on the levels of EGFR.

**Fig 2.**
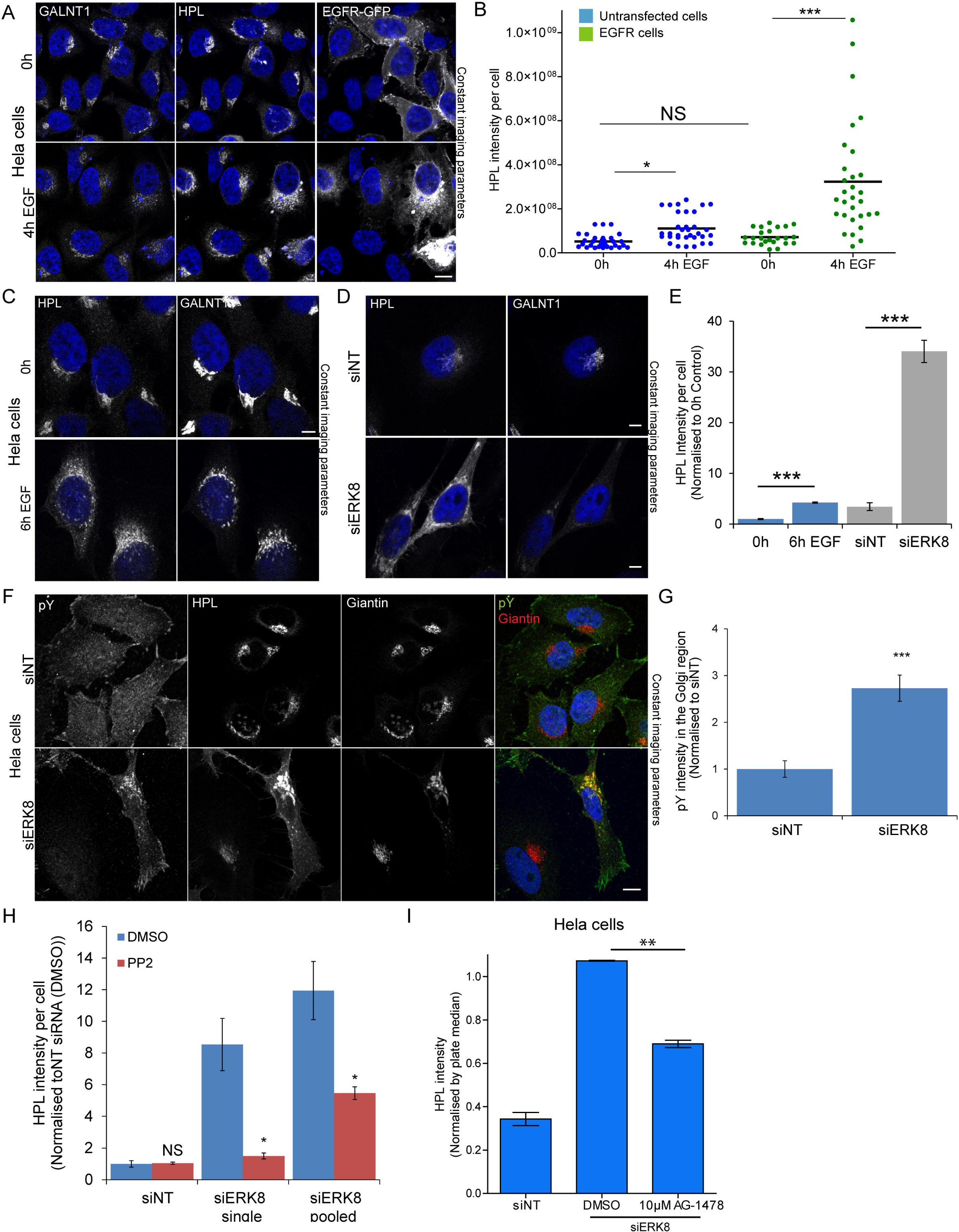
Drivers of GALA: EGFR expression and ERK8 depletion induce more acute GALA activation. (A) Cells transfected with EGFR-GFP stimulated with 100 ng/ml EGF for 4 hours (60X magnification). Scale bar: 10 μm. (B) Quantification of HPL intensity per cell that were untransfected (blue) or transfected with EGFR-GFP (green) from two independent experiments. At least 30 cells were quantified for each condition. Statistical significance (p) measured by two-tailed paired t test. *, p < 0.05 and ***, p < 0.001 relative to mean HPL staining in unstimulated cells (0 h). NS, not significant. Representative images of Hela cells stimulated with (C) 100 ng/ml EGF for 6 hours (D) ERK8 knockdown (“siERK8”) and stained with HPL and GALNT1. Images were acquired at 100x magnification under immersol oil. Scale bar: 10 μm. (E) Quantification of HPL intensity of EGF stimulated (blue) and ERK8 depleted cells (grey) using ImageJ analysis. ***, p < 0.001 relative to mean HPL staining in corresponding unstimulated cells (0h) or non-targeting siRNA (“siNT”) cells. (F) Phosphotyrosine staining (pY20) of Hela cells treated with“siNT” or siERK8” (100X magnification). Scale bar: 10 μm. (G) Quantification of pY20 staining intensity in the Golgi region of siNT or siERK8 treated cells using the ‘Translocation-enhanced’ module on MetaXpress. Images (20X magnification) were acquired on IXM. Statistical significance (p) measured by two-tailed paired t test.***, p < 0.001 relative to siNT cells. (H) Quantification of HPL intensity of siERK8 treated with DMSO control or 10 μM Src family kinase inhibitor PP2 for 24 hours (See images in S2B). “siERK8 single” refers to single siRNA while “siERK8 pooled” refers to a pool of 4 different siRNAs. Statistical significance (p) measured by two-tailed paired t test. ***, p < 0.001 relative to siNT-DMSO control. (I) HPL staining intensity of siERK8 treated with 10 μM EGFR inihibitor AG-1478 or DMSO control. Statistical significance (p) measured by two-tailed paired t test. **p<0.01 relative to siERK8-DMSO control cells. (see images in Fig S2C)

### ERK8 depletion induces a marked relocation of GALNTs

EGF stimulation effect on the intracellular distribution of GALNTs requires optimal conditions of imaging. In many cases, GALNTs staining in the ER is weak and difficult to detect (Fig 2C). This is a consequence of the high dispersion effect of relocation. The ER is a much larger organelle than the Golgi and is distributed over the whole cytoplasm, so the GALNT signal is highly diluted. Because Tn is a more abundant antigen and its levels increase after relocation, its staining can be more easily detected in the ER (Fig 2C). We find that, albeit it is indirect, measuring total levels of Tn is a more reliable and sensitive approach to quantify relocation.

We previously reported that ERK8 depletion induces the redistribution of GALNTs to the ER, quantified via measurement of colocalization with the ER marker Calreticulin [20]. Here, we directly compared EGF stimulation effects with ERK8 depletion (Fig 2C, 2D, 2E). While EGF stimulation induces a 2-fold increase in Tn staining, ERK8 depletion increased signal by 10-fold (Fig 2E). This difference in magnitude is reflected at the Tn and GALNTs level. Upon ERK8 depletion, there is a clear decrease of GALNT1 at the Golgi after ERK8 depletion (Fig 2D). We verified by western blot that GALNT1 protein levels remain constant (Fig S2A). Thus the decrease observed is due to the protein dispersion in the ER.

### GALA activation by ERK8 depletion involves EGFR signaling

We previously reported that growth factor stimulation leads to displacement of ERK8 from Golgi membranes [20]. Here, we tested if ERK8 depletion leads to activation of a tyrosine kinase at the Golgi level. We labeled ERK8 depleted cells with a phospho-tyrosine antibody, PY20 (Fig 2F). Phosphotyrosine staining increased at the Golgi level by about 3-fold (Fig 2G). This is highly reminiscent of the effect of Src activation at the Golgi [25]. Furthermore, treating cells with the Src family tyrosine kinase inhibitor PP2 significantly reduced Tn levels (Fig 2H, S2B). This data indicates that a tyrosine kinase has been activated upon ERK8 depletion. We next treated ERK8 depleted cells with the EGFR inhibitor AG-1478 and found that it reduced Tn levels by at least 40% (Fig 2I, S2C). This result indicates that EGFR signaling is at least partially required to mediate the effects of ERK8 depletion.

We finally tested how general is ERK8 negative regulation of GALA, we repeated the knockdown in two other cell lines. SKOV3 is a cell line originating from an ovarian cancer, with epithelial characteristics. ERK8 depletion led to an 8 fold increase in Tn levels in these cells (Fig S2D, S2E). By contrast, knockdown in HepG2, an hepatocellular carcinoma, did not induce significant GALA. Thus ERK8 negative regulation of GALA is cell type dependent.

### Tn levels increase after EGF stimulation are due to GALNT1/2 activity

HPL and VVL stainings have been associated with structures alternative to Tn. A concern of Dr. Tabak during our exchange of emails was that we were using HPL or VVL staining instead of relying only on GALNTs. We thus performed a double knock-down of GALNT1 and 2, the main GALNTs expressed in Hela cells [24]. Depleting them completely abrogated the increase in Tn induced by EGF (Fig 3A, 3B).

**Fig 3.**
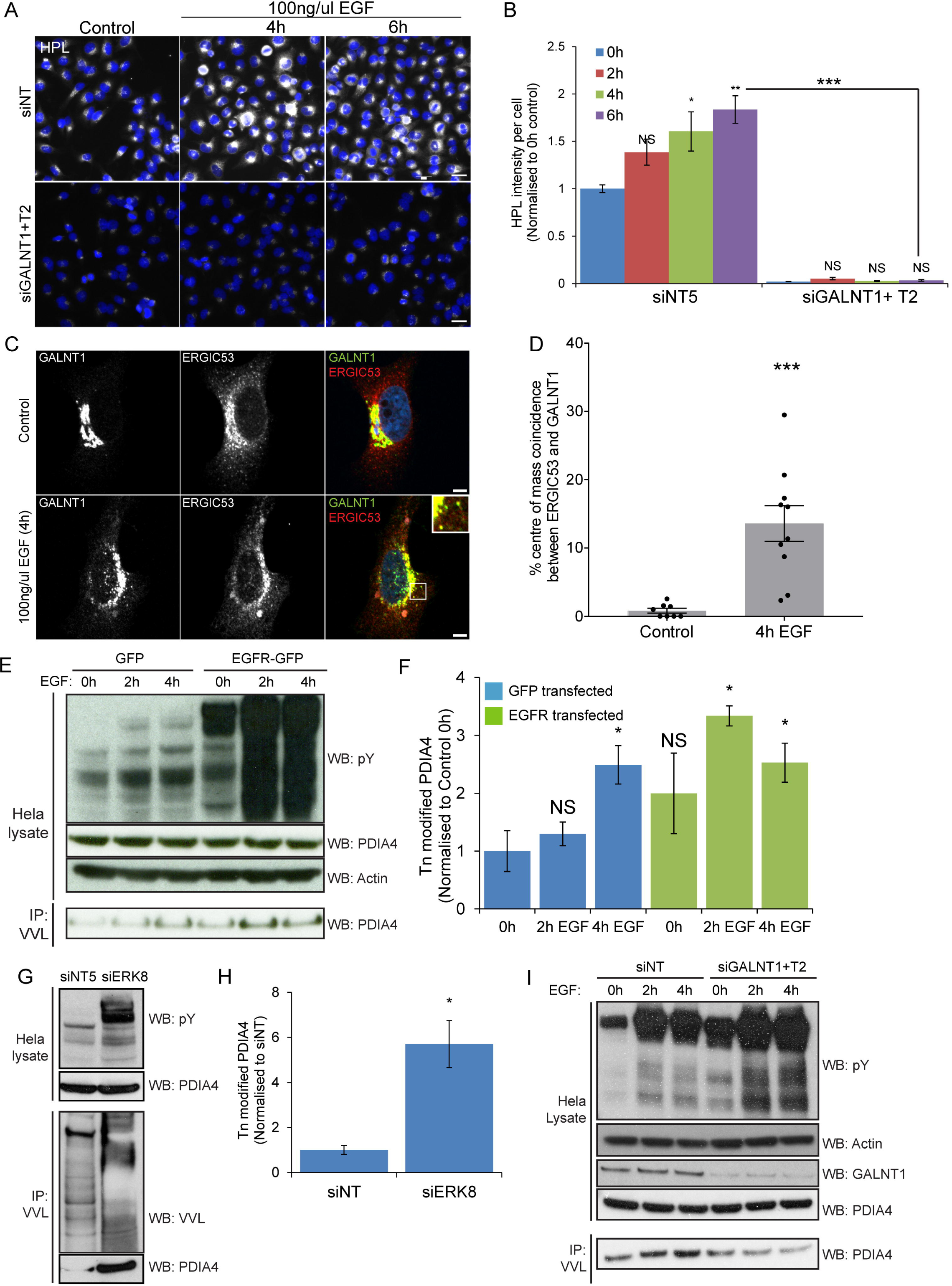
Markers of GALA: O-glycosylation ER resident protein PDIA4 is a reliable marker of GALA activation. (A) HPL staining of GALNT1 and GALNT2 siRNA (“siGALNT1+T2”) and siNT treated cells over time of EGF stimulation. Scale bar: 30 μm. (B) Quantification of HPL intensity in (A). Statistical significance (p) measured by two-tailed paired t test.*, p < 0.05, **, p<0.01 and ***, p < 0.001 relative to unstimulated cells (0h). NS, not significant. (C) GALNT1 and ERGIC53 staining in EGFR expressing cells unstimulated or stimulated with 100 ng/ml EGF for 4 hours (100X magnification). Scale bar: 5 μm. (D) Quantification of the percentage of ERGIC53 intensity centres colocalizing with GALNT1 centres. Statistical significance (p) measured by two-tailed paired t test.***, p < 0.001 relative to untreated cells. (E) Immunoblot analysis of VVL IP on Hela cells expressing EGFR or GFP control that were stimulated with 100 ng/ml EGF over the indicated durations. (F) Levels of Tn modified endogenous ER resident PDIA4 in EGFR or GFP expressing cells over time of EGF stimulation in (EC) from 2 independent experiments. (G) Immunoblot analysis of the levels of Tn modified PDIA4 in Hela cells treated with siNT or siERK8. (H) Levels of Tn modified endogenous ER resident PDIA4 in (G) from 3 independent experiments. (I) Immunoblot analysis of VVL IP on Hela EGFR cells treated with siNT or siGALNT1+T2 which were stimulated with 100 ng/ml EGF over the indicated durations.

### GALA effect can be quantified through punctate structures where ERGIC53 and GALNTs colocalise

As reported in 2010, GALNT staining reveals increased punctate structures at the periphery of the Golgi after 4 hours of EGF stimulation [4]. These structures partially colocalized with ER-to-Golgi intermediate compartment (ERGIC) marker ERGIC53. Colocalization of GALNT and ERGIC53 could thus in theory be used to measure GALA activation. As the two markers appear to overlap only in a fraction of vesicular structures, we employed object-based colocalization analysis that considers each subcellular object as a unique structure. Traditional colocalization methods with Pearson’s or Mandel’s correlation analysis considers global estimation of the entire image and hence would not be informative in this case [26]. By setting a fixed threshold to distinguish objects from noise and background in each staining, the intensity centres of each subcellular structure is determined. A particular structure shows colocalisation if the distance between the centres is less than the imaging resolution. The percentage of structures colocalising in both GALNT and ERGIC53 images was calculated. Using this analysis method, we observed about 14-fold increase in the the population of ERGIC53 structures colocalizing with GALNT1 at four hours post EGF treatment in cells expressing EGFR (Fig 3C, 3D).

### The ER resident protein PDIA4 is a reliable marker of GALA activation

As discussed above, microscopy imaging to quantify GALA can present certain technical limitations. However, the relocation of GALNTs to the ER also induces measurable biochemical changes. We recently identified PDIA4 as a resident ER protein that gets hyperglycosylated upon GALA activation. PDIA4 is hyperglycosylated in tumors that show increased levels of Tn [19].

When Hela cells were stimulated with EGF, we found that PDIA4 glycosylation increased between two and three fold. After transient over-expression of EGFR and stimulation by EGF, it was stimulated by nearly 4-fold (Fig 3E,3F). We also measured PDIA4 glycosylation upon ERK8 depletion in HeLa cells and, consistent with our previous observations, obtained a robust, ~6-fold increase in levels (Fig 3G, 3H). To demonstrate that this change in the glycosylation status of PDIA4 is indeed dependent on GALNTs, we proceeded to deplete GALNT1 and GALNT2 with siRNA. We found that the increased O-glycosylation of PDIA4 in EGF stimulated cells was abolished (Fig 3I). This set of results indicate that PDIA4 is hyper-glycosylated after growth factor stimulation, confirming the relocation of GALNTs from Golgi to ER.

## Discussion

Factors influencing the reproducibility of molecular biological studies are unfortunately varied and numerous. Even cell lines in culture are complex and variable systems with genetic drift and components poorly defined such as FBS. In this study, we have sought to understand why Tabak and colleagues claim that they were not able to reproduce our findings published in 2010. We identified two answers. One is linked to a potential source of variability in these experiments: FBS batches. We found that the batch of FBS used to grow cells can significantly affect GALA response. Effects of FBS batch on signaling are well documented and unfortunately hard to control [27]. Herbomel controlled for EGF effect by probing for ERK1/2 activation, not for total tyrosine phosphorylation nor Src activation. It is thus possible that they used cell culture conditions unfavorable for Src activation, which is a key driver of GALA. As they did not test for this, it is not possible to conclude with certainty.

The second answer is less hypothetical and more surprising: Herbomel et al. did in fact obtain relocation of the GALNTs in their experiments. Using the original raw images shared by them, we found that by changing the display method, the dispersion of GALNTs becomes clearly visible. Furthermore, we found that the image analysis protocol they followed would never have allowed detection of the relocation. Adjusting this protocol with the correct parameters, we were able to quantify GALA effect on GALNT1, GALNT2 and Tn staining in the images they acquired.

Scientists often bring their own preconceived ideas to the bench. It is possible that part of the confusion in Tabak’s group arose from the fact that GALA is not an on/off system, but rather a rheostat-type regulatory process. This is in contrast with other signaling events such as activation of ERK1/2, which tends to function rather as an on/off switch. In cells stimulated by growth factors, GALA activation can be relatively moderate, especially in contrast with the situation in tumor cells in situ. GALA levels tend to correlate with the levels of activation of tyrosine kinases as evaluated by phospho-tyrosine levels. Src (or related tyrosine kinases) activation after EGF stimulation tends to be weaker than ERK1/2 (see for instance [28]). GALA response also varies between cell lines. For instance, HEK293T responded with marked increase in phospho-tyrosines as detected by PY20 antibody as well as increase in Tn levels and Tn modification of PDIA4. By comparison, HeLa response was weaker both at the PY20 and PDIA4 Tn levels. As with FBS conditions, controlling for tyrosine kinases (Src and EGFR) activation levels is critical to understand the levels of GALA.

An active form of Src strongly induce the GALNTs relocation process, an observation that could be reproduced by Tabak’s group (e-mail communication). A complication here is that too strong or too long a Src activation can also lead to Golgi fragmentation, especially in HeLa and other epithelial derived cell lines, as previously reported [29]. This can potentially muddle the measurement of relocation. For unknown reasons, Src activation induces more limited fragmentation in fibroblasts cell lines such as WI-38 or mouse embryonic fibroblasts such as the SYF cell line. In cells that express high levels of Src, a very clear segregation of GALNTs from other enzymes is then clearly observable [4].

By comparison with the growth factor stimulation experiment, our data indicate that GALA is more strongly activated in most human breast and liver tumors. Obviously, this is the relevant context in which to consider this pathway. Are growth factors important *in vivo*? This is not clear at this stage. High levels of EGFR expression and activity are known to be associated with many different tumors, including breast [30] and liver [31]; so this receptor could potentially drive relocation in the pathological context. Src activity is strongly activated with many malignant transformation, so this could be a frequent (but not exclusive) driver of GALA in tumors [32]. In fine, most tumors display much higher levels of Tn than normal tissues and we propose that it is due most of the time to significant GALA activation. Now, this claim can be relatively easily assessed by biochemical analysis of the glycosylation status of the ER resident protein PDIA4.

In a world increasingly concerned with scientific reproducibility, it is equally important to recognize the challenges when attempting reproduction [33]. The complexity of biological systems make a full control of experimental parameters difficult while modern cell biology techniques can be challenging to master. In our own group, genome editing by CAS9-CRISPR was not ‘reproducible’ for months. Hopefully, the clarifications provided in this study will help other researchers who wish to study the GALA pathway to address its significance in tumor progression.

## Materials and methods

### Cell culture

Hela cells were obtained from ATCC. Skov-3 cells were a gift from E. Chapeau. Both Hela and Skov-3 cells were grown in DMEM supplemented with 10% fetal bovine serum (FBS). HEK293T cells were a gift from W. Hong (IMCB, Singapore) and were grown in DMEM supplemented with 15% FBS. All cells were grown at 37°C in a 10% CO_2_ incubator.

### Reagents used in study

Recombinant human epidermal growth factor EGF (#SRP3027) was purchased from Sigma aldrich. PDGF-BB Recombinant Human Protein (#PHG0046), *Helix pomatia* Lectin A (HPL) conjugated with 647-nm fluorophore (#L32454) and anti-GFP (#A11122) were purchased from Thermo Fisher Scientific. Anti-Giantin (#ab24586), anti-actin (#ab8227), anti-PDIA4 (#ab155800) antibodies were purchased from Abcam. Anti-pY416-Src (#2101), anti-Src (#2109), anti-pY-ERK1/2 (#4377), anti-ERK1/2 (#4695) antibodies were purchased from Cell Signaling Technology, Inc. Biotinylated Vicia Villosa Lectin (VVL; #B-1235) and agarose bound VVL (#AL-1233) were purchased from Vector Laboratories, Inc. Anti-phosphotyrosine clone PY20 (#05-947) and anti-phosphotyrosine clone 4G10 (#05-321) were purchased from Merck Millipore. Fetal bovine serum (FBS1; Gibco #10500-064 and FBS2 Gibco #16140-071) were purchased from Thermo Fisher Scientific.

Mouse anti-GALNT1 antibody in hybridoma supernatant was a gift from H. Clausen (University of Copenhagen, Denmark). GFP tagged epidermal growth factor receptor (EGFR) was a gift from W. Hong. siRNAs were purchased from Dharmacon, Inc.

EGFR inhibitor AG-1478 (#S2728) was purchased from Selleckchem. Src family kinase inhibitor PP2 (#529573) was purchased from Merck Millipore.

### Cell transfection, growth factor stimulation and drug treatments

Plasmid transfection in Hela cells and HEK293T cells were performed with Fugene HD (Promega) and Lipofectamine 3000 (Thermo Fisher Scientific) respectively. Cells were transfected for at least 24 hours before further manipulations. siRNA transfection was described in detail in [34]. Briefly, 5μl of 500nM siRNA was mixed with 0.4μl of Hiperfect (Qiagen, #301705) and 14.6μl of Optimem (per well) for 20 min complexation, followed by the addition of 8000 cells to the transfection mix.

For growth factor stimulation, cells were washed twice using Dulbecco’s phosphate-buffered saline (D-PBS) before overnight serum starvation in serum-free DMEM. Human recombinant EGF (100 ng/ml; Sigma-Aldrich) or mouse recombinant PDGF-bb (50 ng/ml;Invitrogen) were added for various durations before fixation with paraformaldehyde or lysed with RIPA lysis buffer.

For drug treatments, drugs were prepared by reconstituting in DMSO in concentrated stock solutions. The drugs or DMSO control were diluted in media containing 10% FBS and added to cells for 6 hours (AG-1478) or 24 hours (PP2) before fixation.

### Immunofluorescence (IF) microscopy

For automated imaging, Hela cells were seeded in a 96-well clear and flat-bottomed black imaging plate (Falcon, #353219) and incubated overnight at 37°C and 10% CO_2_ before growth factor stimulation or plasmid transfection. IF staining procedures was performed as reported in [4,35]. Cells were washed once with D-PBS before fixation with 4%-parafomaldehyde −4% sucrose in D-PBS for 10 minutes. Cells were washed once with D-PBS and permeabilised with 0.2% Triton-X for 10 minutes before the addition of primary antibody in blocking buffer 2% FBS in D-PBS. After primary antibody incubation, cells were washed with blocking buffer for 3 times and subsequently stained for 20 minutes with 2 μg/ml HPL, 5 μg/ml secondary Alexa Fluor–conjugated antibody and 1 μg/ml Hoechst (Invitrogen) in blocking buffer. Cells were then washed for 3 times with D-PBS before imaging. During automated image acquisition, nine sites per well were acquired sequentially with a 20X magnification Plan Fluor on automated imaging system (ImageXpress MICRO [IXM], Molecular devices, LLC). At least three replicate wells for each condition was performed.

For high resolution microscopy, cells were seeded onto glass coverslips in 24-well dishes (Nunc, Denmark). The procedures for IF staining were the same as described above. To observe phosphotyrosine staining, the cells were permeabilized with 0.2% Triton-X for 2 hour at room temperature and stained with anti-phosphotyrosine clone PY20 antibody diluted in 2% FBS in D-PBS overnight. Cells were mounted onto glass slides using FluorSave (Merck) and imaged with immersol oil using an inverted confocal microscope (Zeiss LSM800) at 40X, 60X or 100X magnification.

### Quantification of Immunofluorescence (IF) Staining

Image analysis of HPL intensity of images acquired on IXM was performed using MetaXpress software (version 5.3.0.5). For each well, total HPL staining intensity and nuclei number was quantified using the ‘Transfluor HT’ application module in the software. Hundreds of cells from at least three wells per experiment were quantified. Two experimental replicates was performed.

Image analysis of phosphotyrosine levels at the Golgi were quantified using the ‘Translocation-enhanced’ application module in the software whereby the intensity of PY20 staining within the Golgi area (demarcated by Giantin staining) per cell was quantified. Hundreds of cells from at least three wells per experiment were quantified. Two experimental replicates was performed.

For ImageJ analysis of HPL intensities, images were exported as 16-bit TIFF files. A fixed threshold for a mask on the HPL channel was set and the total intensity per image was calculated based on the total pixel count multiplied by the intensity per pixel. The HPL intensity per cell was then obtained by normalising the total image intensity with the nuclei count.

For analysis of Mander’s coefficient, the images were analysed with ImageJ plugin (Just Another Colocalization Plugin, JACOP). A fixed threshold to distinguish from background signal was set on both HPL and Golgi channels for all images and the corresponding fraction of HPL coincident with Golgi channel (“M1”) and fraction of Golgi coincident with HPL (“M2”) was then used to quantify the colocalization between the markers.

Tabak’s group provided one representative image stack from each experimental condition in .nd2 file format. Each image stack comprise of a z-stack of 61 images for each channel. Each image contains 3-4 cells acquired at 60x magnification. To analyse images provided by Tabak’s group, an ImageJ plugin (Nikon ND2 Reader) was used to export z-stacks of each channel in .avi format. The Mander’s coefficient was analysed with JACOP using the z-stacks from different channels whereby a fixed threshold was set for each of the two channels in the set of images within the same experiment. The maximum projection of the z-stack was presented in the figures.

Image analysis of phosphotyrosine levels at the Golgi were quantified using the ‘Translocation-enhanced’ application module in the software whereby the intensity of PY20 staining within the Golgi area (demarcated by Giantin staining) per cell was quantified.

For object-based colocalization analysis of ERGIC53 and GALNT1 punctate structures, a fixed threshold for each of the channels was set to distinguish discrete objects and from background noise. The intensity centres of each discrete objects i.e. centroids of each channel was calculated in the ImageJ plugin (JACOP). Two objects were considered to colocalise if the distance between their centroids was less than the resolution of the microscope used and the percentage of the objects in the ERGIC53 channel colocalizing with objects in the GALNT1 channel was calculated for each cell.

### Lectin immunoprecipitation (IP)

Cells were washed twice with ice-cold D-PBS before lysis with RIPA lysis buffer containing protease and phosphatase inhibitor (Roche). The protein concentrations of clarified cell/ tissue lysates were measured using the Bradford reagent (Bio-Rad) and normalised across samples. At least 1mg of total lysate was incubated with VVL-conjugated beads for overnight at 4°C. The beads were washed at least three times with RIPA lysis buffer, before the precipitated proteins were eluted in 2x LDS sample buffer with 50mM DTT by boiling at 95°C for 10 minutes. The samples were resolved by SDS-PAGE electrophoresis using bis-tris NuPage gels as per the manufacturer’s instructions (Thermo Fisher Scientific) and transferred to nitrocellulose membranes. Membranes were then blocked using 3% BSA dissolved in Tris buffered saline with tween (TBST: 50 mM Tris [pH 8.0, 4°C], 150 mM NaCl, and 0.1% Tween 20) for 2 hours at room temperature before incubation with lectin or antibodies overnight. Membranes were washed at least three times with TBST before incubation with secondary HRP-conjugated antibodies (GE Healthcare). Membranes were further washed at least three with TBST before ECL (GE Healthcare) exposure.

## Supplemental figures

**Fig S1. Tn staining and GALNT2 analysis in growth factor stimulated cells from Bard’s lab and Tabak’s lab**.

(A) HPL staining of cells stimulated with EGF over the indicated durations performed in Bard’s lab. The Golgi is demarcated by MannII-GFP. Scale bar: 30 μm. (B) Mander’s coefficient to quantify the level of colocalization between HPL and Golgi marker MannII-GFP over time of EGF stimulation in (A). M1 represents the fraction of HPL staining overlapping the Golgi and M2 represents the fraction of Golgi overlapping HPL staining. Values were normalised with respect to unstimulated control cells (0 h). Hundreds of cells were quantified. Statistical significance (p) measured by two-tailed paired t test. **, p < 0.001 relative to unstimulated cells (0 h). (C) Quantification of Mander’s coefficient of GALNT2 and Golgi marker TGN in representative images provided by Tabak’s group. (D) Maximum projection from representative images stained with HPL and ER marker CANX provided by Herbomel et al. Scale bar: 5 μm. (E) Maximum projection images of ER marker CANX in Fig 1F. Scale bar: 5 μm

**Fig S2. ERK8 depletion does not affect GALNT protein levels and occurs through EGFR pathway**.

(A) Immunoblot analysis of GALNT1 levels in Hela cells depleted with ERK8 single (“siERK8 (single)) or ERK8 pooled (“siERK8 (pooled)) siRNA. (B) HPL staining of ERK8 depleted Hela cells treated with DMSO control, 10 μM Src inhibitor PP2 or 10 μM Src Kinase Inhibitor I (SKI-I) for 24 hours. Scale bar: 30 μm (C) HPL staining of ERK8 depleted Hela cells (“siERK8”) treated with 10 μM EGFR inhibitor AG-1478 or DMSO control. Scale bar: 30 μm. (D) HPL staining of ERK8 depleted Skov-3 cells. Scale bar: 30 μm. (E) Quantification of HPL intensity in (D). Statistical significance (p) measured by two-tailed paired t test.*, p < 0.05 relative to siNT control.

